# A modular dCas9-SunTag DNMT3A epigenome editing system overcomes pervasive off-target activity of direct fusion dCas9-DNMT3A constructs

**DOI:** 10.1101/266130

**Authors:** Christian Pflueger, Dennis Tan, Tessa Swain, Trung Nguyen, Jahnvi Pflueger, Christian Nefzger, Jose M. Polo, Ethan Ford, Ryan Lister

**Author notes:** Equal contribution (Christian Pflueger, Dennis Tan). Correspondence (R.L.) and (E.F.).

## Abstract

DNA methylation is a covalent modification of the genome that plays important roles in genome regulation and vertebrate development. Although detection of this modification in the genome has been possible for several decades, the ability to deliberately and specifically manipulate local DNA methylation states in the genome has been extremely limited. Consequently, this has impeded the direct determination of the consequence of DNA methylation on transcriptional regulation and transcription factor binding in the native chromatin context. Thus, highly specific targeted epigenome editing tools are needed to address this outstanding question. Recent adaptations of genome editing technologies, such as the fusion of the DNMT3A methyltransferase catalytic domain to catalytically inactive Cas9 (dC9-D3A), have aimed to provide new tools for altering DNA methylation at desired loci. Here, we performed a deeper analysis of the performance of these tools, revealing consistent off-target binding events and DNA methylation deposition in the genome, limiting the capacity of these tools to unambiguously assess the functional consequences of DNA methylation. To address this, we developed a modular dCas9-SunTag (dC9Sun-D3A) system that can recruit multiple DNMT3A catalytic domains to a target site for editing DNA-methylation. dC9Sun-D3A is tunable, specific and exhibits much higher induction of DNA methylation at target sites than the dC9-D3A direct fusion protein. Importantly, genome-wide characterization of dC9Sun-D3A binding sites and DNA methylation revealed minimal off-target protein binding and induction of DNA methylation with dC9Sun-D3A, compared to pervasive off-target binding and methylation by the dC9-D3A direct fusion construct. Furthermore, we used dC9Sun-D3A to test the impact of DNA methylation upon the DNA binding of CTCF and NRF1 upon targeted methylation of their core binding sites, demonstrating the binding sensitivity of these proteins to DNA methylation *in situ*. Overall, this modular dC9Sun-D3A system enables precise DNA methylation deposition with the lowest amount of off-target DNA methylation reported to date, allowing accurate functional determination of the role of DNA methylation at single loci.

## Introduction

DNA methylation has been shown to play a critical role in development, pathogenesis of various disease states, and is frequently associated with transcriptional repression (Okano et al. 1999; Jackson-Grusby et al. 2001; Egger et al. 2004; Smith & Meissner 2013; Bernstein et al. 2007; Perino & Veenstra 2016; Li et al. 1992; Gao & Teschendorff 2017). In recent years, single base resolution methylome maps have been generated for different cell types and organisms, providing insights into the many possible functions of DNA methylation in the genome (Cokus et al. 2008; Lister et al. 2009; Irizarry et al. 2009; Maunakea et al. 2010; Stadler et al. 2011; Lister et al. 2013). Although comparative analyses of DNA methylation patterns with other genomic data such as transcription factor binding, gene expression and chromatin state have been used to infer the functions of this DNA modification, these techniques only provide correlative information. Therefore, a growing body of research is aimed at disentangling the cause or consequence of gene repression by DNA methylation, including its potential role in the shaping of the transcription factor (TF) binding landscape.

Traditional approaches to directly assess the relationship between CG methylation (mCG) within TF binding motifs and the occupancy of potentially mCG-sensitive TFs has relied on either the artificial insertion of DNA sequences that contain a TF binding motif with differing mCG states, biochemical experiments leveraging gel shift properties, pharmacological inhibition, or genetic perturbation of DNA methyltransferases (DNMTs) (Renda et al. 2007; Stadler et al. 2011; Maurano et al. 2015). The use of DNMT inhibitors, such as 5-azacytidine, were previously used to alter DNA methylation states and infer the influence upon TF binding (Wang et al. 2012; Maurano et al. 2015). However, these pharmacological inhibitors of DNMTs alter the DNA methylation landscape globally, induce broad transcriptional changes, and result in highly pleiotropic off-target effects. Therefore, these are not suitable to understand the nuanced impact of DNA methylation on TFs at specific binding sites. More recently, a systematic evolution of ligands (methylated and unmethylated short double stranded DNA) by exponential enrichment (SELEX)-based investigation of the effect upon TF binding capacity *in vitro* of mCG in the core DNA binding site for hundreds of TFs and TF binding domains revealed that the majority of TFs are sensitive to mCG, resulting in either increased or reduced DNA binding affinity (Yin et al. 2017). However, *in vitro* binding assays lacking the native chromatin context and the artificial insertion of DNA sequences cannot accurately recapitulate an actual biological process, while pharmacological and genetic perturbations of DNMTs cause undesired global depletion of mCG. Thus, these techniques suffer from confounding secondary effects due to a lack of target selectivity, and are therefore unable to accurately address the role of mCG in regulating mCG-sensitive TF binding. Therefore, there is a need to develop epigenome editing tools that can achieve targeted and highly specific modulation of mCG states at TF binding motifs in order to clarify the roles of mCG in shaping the mCG-sensitive TF occupancy landscape.

A major challenge has been the development of precise, adaptable tools that are capable of directing targeted DNA methylation to individual loci in different genomic contexts. Advances have been made by directly fusing DNA methyltransferase enzymes (e.g. the *de novo* mammalian DNA methyltransferase 3a (DNMT3A) or the prokaryotic CG methyltransferase, M.SssI/MQ1) to programmable DNA-binding domains including zinc fingers, transcription activator-like effectors (TALEs), and deactivated Cas9 domains (dCas9) to induce targeted deposition of DNA methylation (Rivenbark et al. 2012; Siddique et al. 2013; Nunna et al. 2014; Bernstein et al. 2015; Stolzenburg et al. 2015; Stepper et al. 2017; McDonald et al. 2016; Amabile et al. 2016; Liu et al. 2016; O’Geen et al. 2017; Xiong et al. 2017; Lei et al. 2017; Ford et al. 2017). Thus, in theory, it is now possible to examine context dependent transcriptional changes in response to localised epigenomic changes, but this varies greatly between target sites and DNA-binding domains (Jurkowski et al. 2015; Köferle et al. 2015). Furthermore, we currently understand very little about the off-target effects of these systems upon methylation throughout the genome. Given the importance of target specificity to these systems, gaining a comprehensive understanding of specificity, and developing approaches to improve it, is critical for the implementation and progression of accurate epigenome engineering.

Recently, attempts at investigating TF occupancy and mCG binding sensitivity have focused on the CCCTC-binding factor (CTCF), a transcription factor involved in DNA looping and chromatin architecture (Phillips & Corces 2009; Hashimoto et al. 2017). These studies have utilized direct fusion constructs between dCas9 and DNMT3A (Liu et al. 2016) or M.SssI (Lei et al. 2017). The latter achieved highly specific, but only low level, induction of methylation at a limited number of CG sites across their target region (Lei et al. 2017), while the former attained higher levels of methylation induction but also reported off-target methylation (Liu et al. 2016). This suggests limited versatility and specificity of these single fusion constructs, despite both studies reporting reduced CTCF occupancy upon changes in mCG state. Furthermore, scant attention has been paid to the binding specificity and off-target DNA methylation delivered by these epigenetic editing tools, which may confound the interpretation of results. Hence, it is paramount to study the sensitivity of TF binding to methylated DNA within their native chromatin context at endogenous loci, with the highly precise targeted DNA methylation editing. Here, we describe the development and comparative analysis of dCas9 based epigenome editing systems that recruit the catalytic domain of DNMT3a, with a particular focus on comprehensive assessment and minimization of off-target binding and DNA methylation induction, and utilization of optimized systems for modulating TF binding.

## Results

### Direct Fusion of dCas9 to the DNMT3A catalytic domain results in high off-target DNA methylation

The direct fusion of dCas9 to the catalytic domain of DNMT3A was previously reported and used to induce cytosine methylation at targeted loci (Qi et al. 2013; Vojta et al. 2016; Amabile et al. 2016; Liu et al. 2016; Stepper et al. 2017). However, the potential off-target binding and methylation induced by these constructs has not been comprehensively assessed. In order to better understand the on-target and off-target effects of this system, we generated two dCas9-DNMT3A constructs, dC9-D3A and dC9-D3A-high (Fig. 1a), where the sole difference was in their puromycin selection marker. While identical in their structural design, dC9-D3A-high had the puromycin N-acetyltransferase fused to a self-cleavable peptide (P2A)(Kim et al. 2011) whereas the dC9-D3A construct had the puromycin selectable gene expressed via its own constitutive promoter. MCF-7 cells or HeLa cells were transiently transfected with these two constructs and a gRNA targeting the UNC5C promoter and incubated for 48 h to allow for adequate protein expression followed by 48 h puromycin selection. Cells were then harvested and DNA or chromatin was extracted for targeted bisulfite sequencing (targeted bsPCR-seq), chromatin immunoprecipitation (ChIP) or ChIP bisulfite sequencing (ChIP-bs-seq) (Brinkman et al. 2012) (Fig. 1b). Initially we measured induced on-target DNA methylation at the UNC5C promoter and a proxy for potential off-target DNA methylation at the promoters of the BCL3 and DACH1 genes (Fig. 1c) in HeLa cells with dC9-D3A or dC9-D3A-high. Surprisingly, dC9-D3A exhibited poor on-target DNA methylation induction at the *UNC5C* promoter, with the highest observed increase in methylation at any single CG site within the region of 21%, and an average ΔmCG increase of 5% over all 62 CGs in the region. In contrast, dC9-D3A-high induced DNA methylation up to 52% at singe CG sites in the *UNC5C* promoter and an average ΔmCG increase of 16% over the promoter region. However, dC9-D3A-high displayed strong off-target DNA methylation activity compared to dC9-D3A at the *BCL3* promoter (average increase of 3% and 0.8%, respectively, over 46 CGs), but less so at the *DACH-1* promoter (average increase of 1.6% and 1.4%, respectively, over 48 CGs)(Fig. 1c). We speculated that the different on and off-target DNA methylation rate was the result of differing protein levels of dC9-D3A and dC9-D3A-high. Indeed, western blot on protein extractions from cells before and after puromycin selection demonstrated that protein levels were substantially higher in cells transfected with dC9-D3A-high compared to dC9-D3A, regardless of selection (Fig. 1d). We consequently reasoned that modulating the on and off-target efficiency for targeted DNA methylation could not be accomplished by merely changing expression levels of the direct fusion constructs, and that there was an inherent tradeoff in the on-target methylation efficacy versus off-target methylation with the direct fusion system. A redesigned, modular system for independent recruitment of the DNMT3A effector may be effective for overcoming this shortcoming (Fig. 1a).

**Figure 1:**
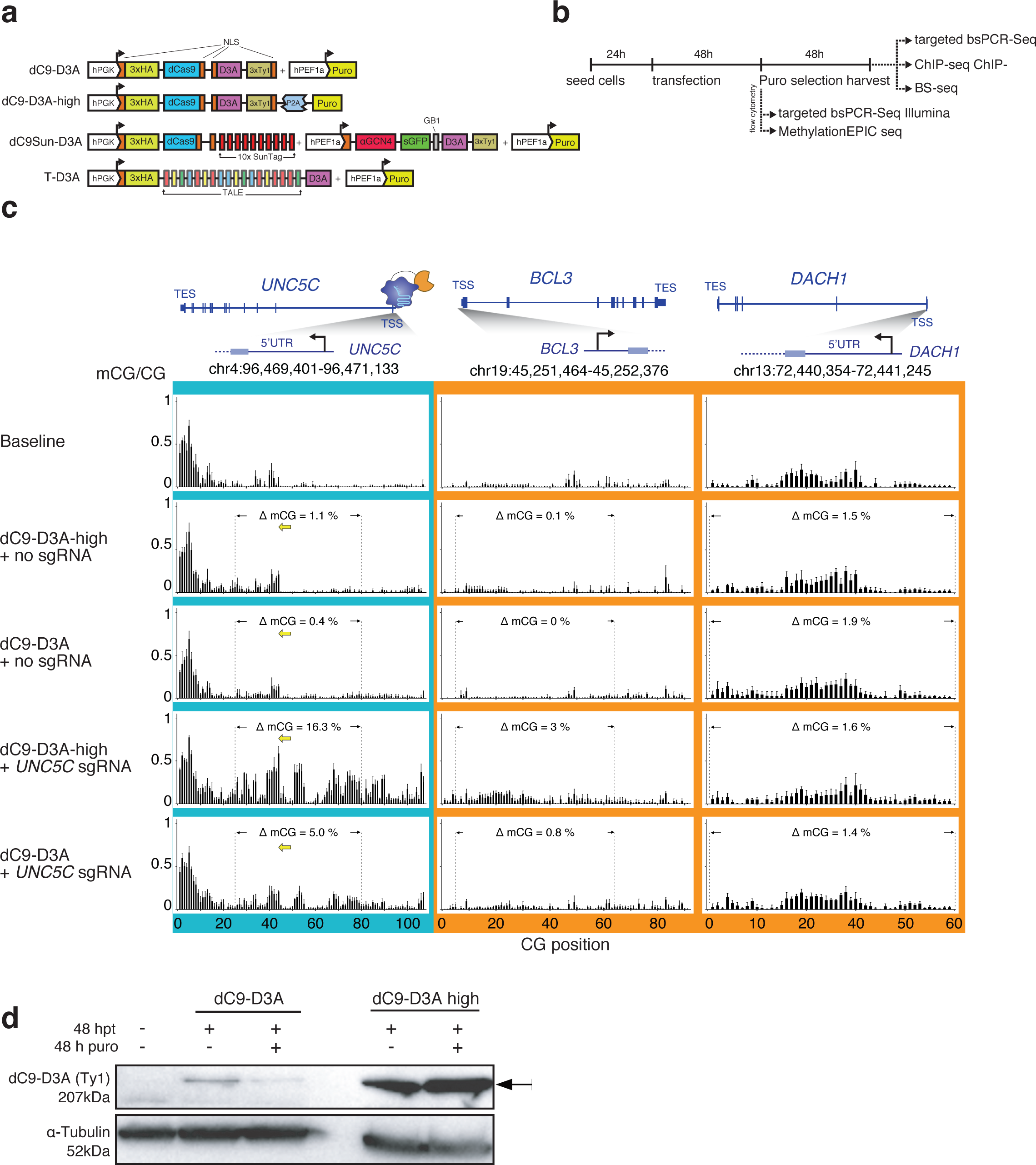
Characterizing on-target and off-target mCG deposition efficiency by dC9-D3A direct fusion system. **(a)** Schematics of the dCas9 (dC9) and TALE (T) constructs used, indicating positioning of nuclear localization sequences (NLS), protein tags (human influenza hemagglutinin - 3xHA, 3xTy1), promoter choice (glycerol kinase promoter - hPGK, human elongation factor-1 alpha promoter - hPEF1a), solubility tag (protein G B1 domain - GB1), selectable marker (puromycin resistance - puro), single-chain Fv antibody against GCN4 domain (αGCN4) and human DNTM3A catalytic domain (D3A). **(b)** Timeline and outline for experimental design for measuring DNA methylation and TF occupancy (CTCF and NRF1). **(c)** Targeted DNA methylation deposition to *UNC5C* promoter in HeLa cells, measured by bsPCR-seq, sgRNA placement shown with yellow arrows, dotted lines indicate interval for CGs included in quantitation. **(d)** Western blot of relative dC9-D3A protein abundance (anti-Ty1) per 50 µg of total cell lysate, 1st lane untransfected HeLa cells, 2nd and 4th lane: 48 h post transfection (hpt), 3rd and 5th lane: 48 hpt and 48 h puromycin (puro) selection, loading control anti-Tubulin, arrow indicates dC9-D3A.

### Adapting a modular dCas9 system for achieving high and specific DNA methylation

A modular dCas9 system has previously been reported utilizing the SunTag array (Tanenbaum et al. 2014). This repeating array of short repeat peptide sequences is fused to dCas9, thus acting as an epitope docking station that allows multiple proteins (fused to the counterpart single-chain antibody, scFv-GCN4, subsequently referred to as αGCN4) to be recruited to a desired target site. We hypothesized that this system would allow us to independently modulate the expression of the DNMT3A catalytic domain and the dCas9-SunTag, with the aim of limiting dCas9-SunTag abundance to favor its binding to its highest affinity sites in the genome while restricting the presence of excess DNMT3A in order to avoid its non-specific activity. Such a system should reduce spurious off-target DNA methylation while maintaining high on-target mCG induction, thus improving on the design of the dCas9 molecule directly fused to a single DNMT3A. We adapted the previously reported SunTag system by fusing ten GCN4 peptides with a flexible linker to dCas9 (dC9Sun) and the counterpart antibody to DNMT3A (αGCN4-D3A; dC9Sun-D3A, Fig. 1a). The resulting dual constructs were tested in a titration series (Fig. 2a) that aimed to maximise on-target DNA methylation deposition at the *UNC5C* promoter, while minimising off-target DNA methylation induction. The *BCL3* promoter was used as an initial reporter locus for detecting the induction of excessive off-target DNA methylation. The entire set of transfections was carried out in HeLa cells which were transfected with a fixed amount of pGK-dCas9-SunTag plasmid and varying amounts of effector (αGCN4-D3A). The latter was titrated from 0 to 33% of the total amount of DNA transfected. The optimum amount of pEF1a-αGCN4-D3A effector that yielded the highest amount of on-target DNA methylation and least amount of off-target DNA methylation was determined to be 4% of the total amount of DNA transfected. The change in DNA methylation (ΔmCG) was calculated by subtracting the average of all mCG/CG ratios in a defined region (e.g. 62 CGs at the *UNC5C* promoter) from the same CG sites in control treated cells (baseline). Our adapted system was able to achieve high on-target mCG deposition of up to 49% at individual CpG dinucleotides and an average ΔmCG of 12.6% over all 62 CG dinucleotides, while maintaining off-target DNA methylation at only 0.7% (Fig. 2a, green highlight). Hereafter, these optimized ratios of pGK-dCas9-SunTag plasmid to pEF1a-αGCN4-D3A effector plasmid were used for the remainder of the study, and will be collectively referred to as dC9Sun-D3A. The use of the SunTag system to recruit DNMT3A via TALEs to the *DACH1* and *UNC5C* promoters also substantially improved on-target mCG deposition compared to a TALE-DNMT3A single fusion (Supplementary Fig. 1 and 2). However, while TALEs are known for their high specificity, they are time consuming to adapt for multiple target sites and challenging to multiplex, hence we focused on the more rapidly configurable dCas9 system for the remainder of the study.

**Figure 2:**
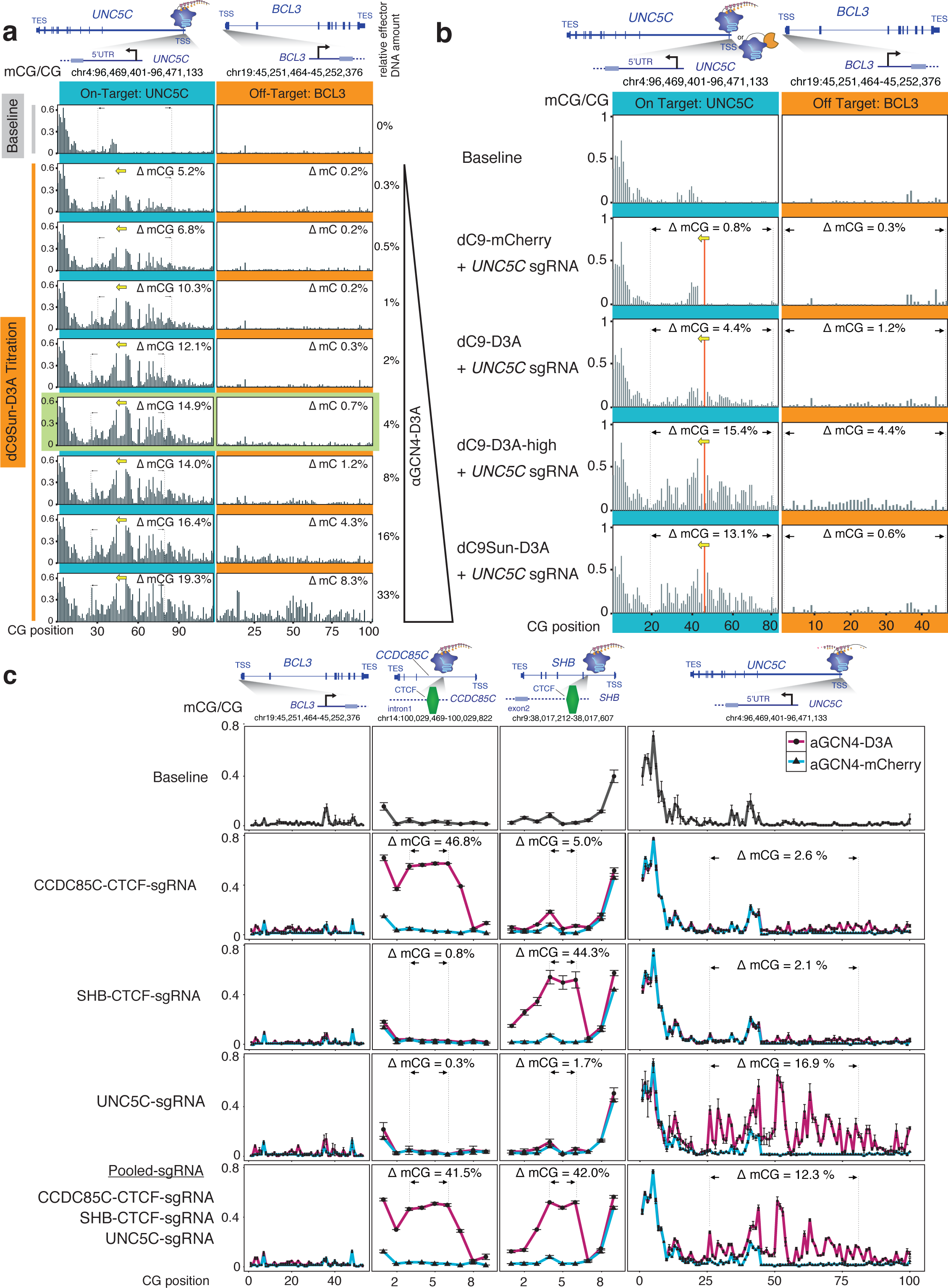
Modular dC9Sun-D3A system outperforms dC9-D3A direct fusion. **(a)** Titration of aGCN4-D3A effector (D3A - human DNMT3A catalytic domain), fraction mCG shown in black bars, dotted lines and black arrows indicate region used to calculate mCG change. **(b)** Comparison of dC9-D3A high, dC9-D3A, dC9Sun-D3A and dC9Sun-mCherry (CRISPRi control) at UNC5C promoter (on-target) vs. BCL3 promoter (off-target) by targeted bsPCR-seq (average mCG/CG, n=3 replicates, SD error bars). **(c)** mCG deposition efficiency by dC9Sun-D3A at three different loci (*CCDC85C, SHB* and *UNC5C* promoter) measured by targeted bsPCR-seq (average mCG/CG, n=3 replicates, SD error bars).

To control for changes in methylation due to the presence of dCas9 binding in the vicinity of our targeted region, we initially used a catalytic mutant of the DNMT3A catalytic

domain (D3AMut), which harboured four alanine substitutions in its catalytic center (F39A, E63A, E155A, R284A). However, when mCG deposition was tested at three target sites (*CCDC85C* intron, *SHB* intron and *MIR152,* Supplementary Fig. 3 a-c) we detected residual mCG deposition of up to 22%, 16% and 8%, respectively. We speculate that the DNMT3A catalytic domain might be able to recruit endogenous functional WT DNMT3A by forming oligomeric complexes as previously described (Holz-Schietinger et al. 2011). Accordingly, to eliminate any potential functional impacts based on residual induction of cytosine methylation, we determined that an αGCN4-mCherry effector would be a more appropriate control for the effect of dCas9 binding alone, compared to the αGCN4-D3AMut construct.

Targeting of the *UNC5C* promoter was repeated with dC9Sun-D3A, dC9-D3A and dC9-D3A-high to compare the performance of each system, measuring the methylation level at CpG dinucleotides surrounding the *UNC5C* target region as well as at the *BCL3* promoter, which served as a proxy for widespread off-target DNA methylation (Fig. 2b). dC9Sun-D3A induced mCG at the *UNC5C* promoter to comparable levels as dC9-D3A-high (ΔmCG over 62 CG sites at *UNC5C* was 4%, 15% and 13% for dC9-D3A, dC9-D3A-high and dC9Sun-D3A, respectively), without the shortcomings of promiscuous off-target methylation (ΔmCG over 46 CG sites at the off-target *BCL3* region was 1.2%, 4.4% and 0.6% for dC9-D3A, dC9-D3A-high and dC9Sun-D3A, respectively). Notably, the off-target mCG induction by dC9Sun-D3A was reduced by more than 80% compared to dC9-D3A-high (Fig. 2b, *BCL3* promoter off-target methylation).

To determine whether this improved DNA methylation induction and reduced off-target are achievable at multiple different loci, the set of target sites was expanded both independently and via multiplexed targeting. This was achieved by selecting a series of CTCF sites that were previously shown to be CG methylation sensitive (Maurano et al. 2015), hypomethylated and occupied by CTCF in HeLa cells. Notably, high mCG induction was consistently achieved across both CTCF core binding sites (*CCDC85C* intron (47%) and *SHB* intron (44%)) and the *UNC5C* promoter (17%) (Fig. 2c, row 2-4). Multiplexing all three single guide RNAs (sgRNAs) resulted in a loss of on-target mCG deposition of less than 5% at all three targets (highlighted regions, Fig. 2c, row 2-4 compared to Fig. 2c, row 5), strongly suggesting that the dC9Sun-D3A system is a viable multiplexing option for mCG deposition. The maximum ΔmCG observed was at the *UNC5C* promoter, with a 57% increase. dC9Sun-D3A was able to induce DNA methylation at distances ≥300 bp, however, the SunTag system did not appear to add any additional distance compared to the dCas9 direct fusion (Supplementary Fig. 4). The same was observed when the T-D3A direct fusion is compared to TSun-D3A, however, adding the SunTag system to a TALE domain targeting the *UNC5C* promoter improves on-target mCG deposition from 30% to 64% (Supplementary Fig. 4).

Furthermore, we investigated whether a similar dC9Sun system was able to facilitate DNA demethylation when coupled to human TET1 (tet methylcytosine dioxygenase 1) catalytic domain (Morita et al. 2016) at the *GAD1* intron #3. We found that the efficacy of the system was highly dependent on the sgRNA placement, with dC9Sun-TET1 achieving up to 60.0% mCG reduction at the *GAD1* intron #3 (Supplementary Fig. 5). Taken together, the system of dC9Sun in combination with epigenetic effectors coupled to αGCN4 prove to be highly efficient in targeted deposition or removal of DNA methylation, without the limitations of off-target effects.

### Global assessment of binding and methylation specificity

In order to more comprehensively assess the specificity of the dC9Sun-D3A system we performed ChIP-seq upon dC9Sun using the 3xHA epitopes present at the N-terminus of the protein, after targeting the construct to intron #1 of *SHB* (Fig. 3a). ChIP-seq peak calling identified 13 significant peaks throughout the entire genome (qvalue < 0.01), with the top 10 most significant peaks shown in Figure 3a. All but one off-target peaks were found to have less than 10% of the ChIP-seq normalized read density compared to the on-target binding site peak in the *SHB* intron (Supplementary Table 1). The reciprocal αGCN4-D3A ChIP-seq experiment using the C-terminal 3xTy1 epitope only yielded only 1 significant peak at the on-target *SHB* site (Supplementary Table 1).

**Figure 3:**
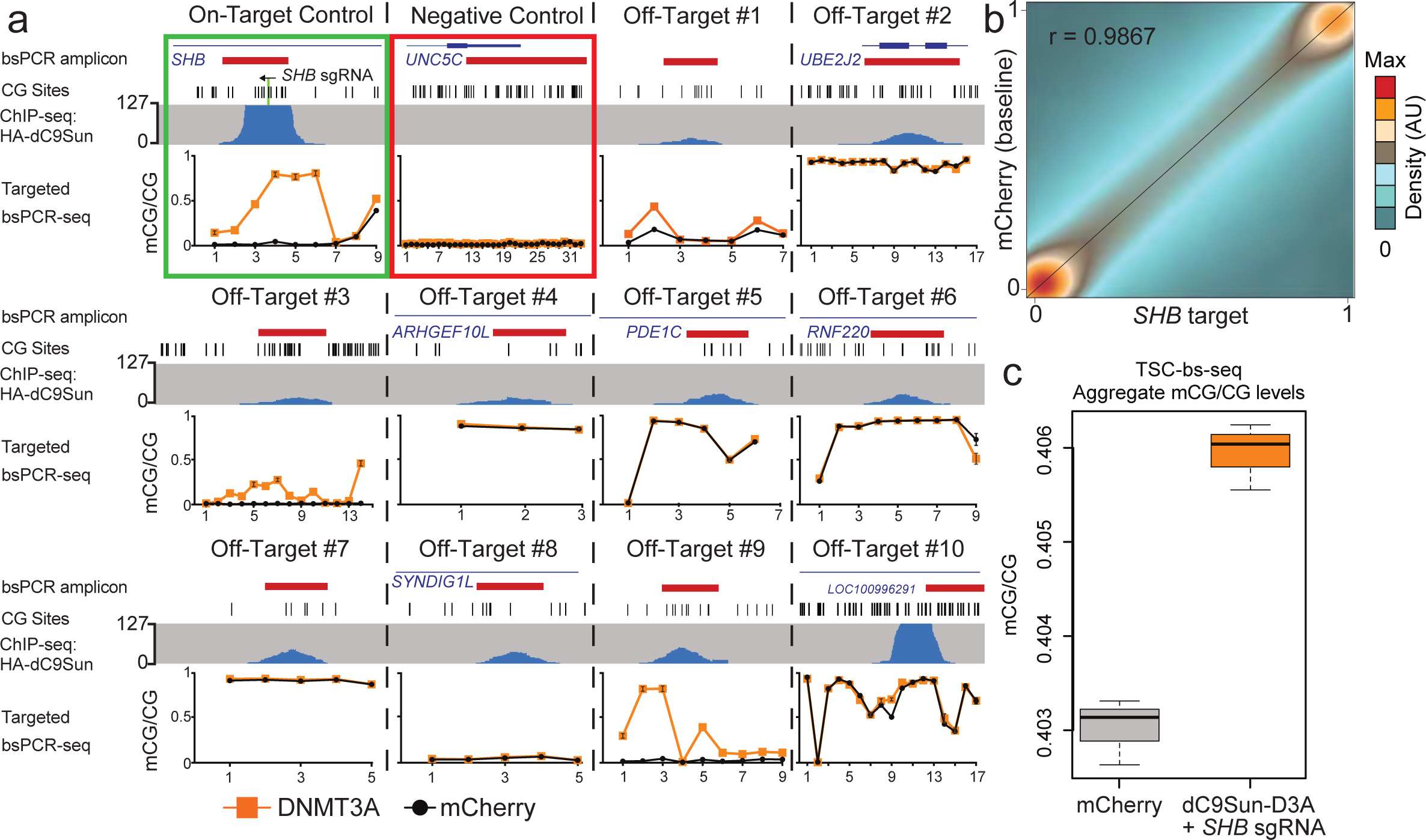
Genome-wide off-target DNA methylation assessment. **(a)** Compilation of dC9Sun-D3A ChIP-seq (blue peaks) and targeted bsPCR-seq for CGs covered by ChIP-seq (bsPCR amplicon location in red), bsPCR-seq mCherry only expressing cells (black line) and *SHB* sgRNA + dC9Sun-D3A (orange line), (sorted cells, n=3 biological replicates, SD error bars). **(b)** Correlation of mCG values for each CG site (> 2.6×10^6^, >=5xcoverage)ofcombinedreplicates(n=3) from lllumina TruSeq Methyl Capture EPIC pulldown experiment, mCherry (baseline) HeLa cells shown on y-axis, *SHB* target dC9Sun-D3A HeLa cells shown on x-axis. Pearson correlation efficent shown in upper left corner. **(c)** Boxplot of mCG/CG from all covered CGs from mCherry only HeLa cells (gray) and *SHB* sgRNA + dC9Sun-D3A (orange) (n=3 biological replicates, median - thick black line, SD error bars).

To further explore the extent of off-target mCG deposition and dC9Sun-D3A occupancy, we designed all three components of the system (dC9Sun, αGCN4-D3A, and *SHB* sgRNA) to express different fluorescent proteins (BFP, GFP and mCherry, respectively), so that cells transfected with all three constructs could be enriched using flow cytometry (Supplementary Fig. 6a). The cell sorting approach, enriching for cells that have an intermediate level of expression of dC9Sun (BFP co-expression) and αGCN4-D3A (GFP fusion), was chosen to firstly guarantee expression of all 3 components (dC9-Sun, aGCN4-D3A and sgRNA) in a single cell, and secondly, to achieve the highest possible on-targeted mCG deposition at *SHB* promoter while keeping off-target mCG change to a minimum (Supplementary Fig. 6b). Next, targeted bsPCR-seq in these sorted cells was performed for off-target sites and the on-target site *SHB* intron. Of all the off-target ChIP peaks, only 3 exhibited an increase in one or more underlying CG dinucleotides of ΔmCG >10% (off-target peak region #1, #3, and #9; Fig. 3a). A plausible explanation for the off-target binding of dC9Sun-D3A to these sites is that they contain partial matches to the *SHB* sgRNA, with the matching bases ranging from 11 to 15 nt out of the 17 nt of the *SHB* sgRNA target sequence; Supplementary Fig. 7). Therefore, the specificity of this system is likely only restricted by the uniqueness of the genomic sequence to which dCas9 is targeted, and could be improved by alternative sgRNA design. Overall, these results demonstrate that the dC9Sun-D3A system can exhibit very high specificity of binding in the genome.

While binding of dC9Sun-D3A appeared to be highly specific, there is the potential that expression of the components of the system could still induce off-target methylation without detectable binding of dC9Sun-D3A. Therefore, to identify any mCG changes more broadly throughout the genome, the Illumina TruSeq Methyl Capture EPIC kit was used for targeted capture and detection of the DNA methylation state at single base resolution (target solution capture bisulfite sequencing, TSC-bs-seq) for 2.6 million CpG sites located in genomic regulatory regions in the DNA isolated from the previously fluorescently sorted cells (Supplementary Fig. 6). This approach allows quantitation of the DNA methylation state of ∼12% of all CGs present in the human genome, which are specifically targeted to cover the majority of known constitutively or conditionally unmethylated and lowly methylated regions in the genome (97% of CpG islands, 95% GenCode promoters, 66% open chromatin regions, 98% FANTOM5 enhancers, 78% transcription factor binding sites). Therefore, this provides an effective approach for high coverage (≥5 reads per CG and replicate) base resolution detection and quantitation of potential off-target methylation in the regions of the genome that could potentially become methylated. These probes captured regions including the on-target *SHB* intron and off-target site #9, where up to 80.5% and 82.1% methylation was observed, respectively, as observed in the targeted bsPCR-seq analysis (Supplementary Fig. 8). Notably, on-target mCG deposition was measured independently to be 80.5% and 78.0% for the CGs in the CTCF binding site (*SHB* intron) by either TSC-bs-seq or targeted bsPCR-seq, respectively. The methylation level quantitated at CG sites by both targeted bsPCR-seq and TSC-bs-seq methods were very similar to each other, with spearman correlation coefficient for controls and dC9Sun-D3A treatment of r=0.976 and r=0.967, respectively (total number of CGs=94, n=3 replicates). Furthermore, both sets of measurements suggest that sorting cells for optimal expression of all three components (dC9Sun, αGCN4-D3A and sgRNA) reduces off-target mCG deposition by 2.1-fold (*UNC5C* promoter) and improves on-target mCG deposition in the cell population by 17.3% compared to puromycin selection (Supplementary Fig. 9). The high specificity of targeted DNA methylation induction by the dC9Sun-D3A system was further demonstrated by the very similar mCG levels (Pearson correlation coefficient r = 0.9867) at the 2,629,232 CGs covered (depth ≥5 reads, median CG coverage ≥16) throughout the targeted regions of the genome between cells expressing the control construct (dC9Sun-mCherry) and those expressing dC9Sun-D3A and the *SHB inton #1* sgRNA (Fig. 3b). This was also observed for all pairwise correlations of each replicate (Supplementary Fig. 10). Moreover, the difference in average fraction mCG/CG for all CGs covered (≥5 reads coverage, 2,629,232 CGs, n=3 replicates) between cells expressing the control construct (dC9Sun-mCherry) and those expressing dC9Sun-D3A and the *SHB intron* sgRNA was less than 0.3% (Fig. 3c). Overall, this demonstrates that the dC9Sun-D3A system exhibits very high specificity in both binding sites and induction of DNA methylation, in stark contrast to the high off-target activity of dC9-D3A (Fig. 1c, 2b), the system that has been most commonly used to date (Amabile et al. 2016; Liu et al. 2016; Vojta et al. 2016).

### Direct assessment of the effect of DNA methylation on DNA-protein interactions

Having established the specificity and efficiency of the dC9Sun-D3A system, we sought to examine the consequences of targeted mCG deposition upon binding of methylation sensitive transcription factors. Previously, Maurano *et al.* (Maurano et al. 2015) reported that a subset of CTCF binding sites are sensitive to mCG reduction, induced by either DNMT triple knockout in HCT116 cells or 5-aza-cytidine treatment in K562 cells. Based on these findings, we speculated that CTCF binding sites sensitive to mCG loss would exhibit a reciprocal phenotype when mCG was specifically deposited in CTCF core binding sites in HeLa cells. To that end, we selected target regions for testing by intersecting the CTCF binding sites that were mCG sensitive in both K562 and HCT116 sites (FDR < 0.01) with CTCF sites occupied in HeLa cells. These sites were further required to each contain at least one hypomethylated CG in their core binding site. Three resulting CTCF binding sites were selected, in 1) a *SHB* intron, 2) a region upstream to *MIR152*, and 3) a *CCDC85C* intron. CTCF binding sites for *MIR152*, *SHB* intron and *CCDC85C* intron are completely unmethylated in HeLa cells, as determined by whole genome bisulfite sequencing (WGBS, Fig. 4a, 4b and Supplementary Fig. 11, HeLa WGBS track). This observation was confirmed by targeted bsPCR-seq, where the amplicon covered the entire CTCF core binding site (Fig. 2c, baseline panel, *CCDC85D* and *SHB*). Next, we targeted mCG to the CTCF core binding sites by recruiting either dC9Sun-D3A or dC9Sun-D3AMut with sgRNAs binding within 100 bp of the CTCF core binding site. dC9Sun-D3AMut was used to control for possible steric hindrance effects on CTCF binding, however we subsequently utilized dC9Sun-mCherry since dC9Sun-D3AMut exhibited residual DNA methylation activity (Supplementary Fig. 3a-c). After 48 h of puromycin selection, the on-target mCG deposited at the CTCF core binding sites were 62% for the single CG in the CTCF binding site in *SHB* (Fig. 4c), 31% and 46% for the two CGs in the CTCF binding site upstream of *MIR152* (Fig. 4d), and 52% and 35% for the two CGs in the CTCF binding site in *CCDC85C* (Supplementary Fig. 11b). We then investigated whether CTCF binding was impacted by the targeted DNA methylation deposited in its core binding site by performing CTCF ChIP-seq in duplicates for each condition. Importantly, all three CTCF binding sites, which were independently targeted by dC9Sun-D3A, showed a significant decrease in CTCF occupancy compared to dC9Sun-D3AMut (0.36-fold, 0.31-fold and 0.54-fold reduction in normalized ChIP-seq read density for *SHB*, *MIR152* and *CCDC85C*, respectively (edgeR, FDR = 4.6×10^−3^, 2.9×10^−5^, 2.7×10^−15^, respectively). Notably, the reduction in CTCF binding at the targeted loci (*MIR152*, *SHB* and *CCDC85C*) were ranked the most or second most decreased CTCF peaks compared to control, as determined by edgeR analysis of normalized peak counts (*SHB* - Fig. 4e, *MIR152* - Fig. 4f, *CCDC85C* - Supplementary Fig. 11c). Further, we performed CTCF ChIP-bisulfite sequencing (ChIP-bsseq)(Hon et al. 2012; Brinkman et al. 2012; Statham et al. 2012) to assess the DNA methylation status of the DNA that was directly bound by CTCF (Fig. 4 a,b and Supplementary Fig. 11a). This confirmed that the DNA bound by the remaining CTCF was methylated, and comparable to the initial on-target DNA methylation measured by targeted bsPCR-seq (Fig. 4c and 4d, top panel comparing gray to colored circles), suggesting that CTCF occupancy is reduced by >30% for all 3 CTCF target sites (*SHB*, *MIR152* and *CCDC85C*) due to mCG deposition. The remaining chromatin bound CTCF was found to have mCG in its core binding, indicating a possible transition state, where CTCF is either poised to leave or mCG is targeted for active DNA demethylation.

**Figure 4.**
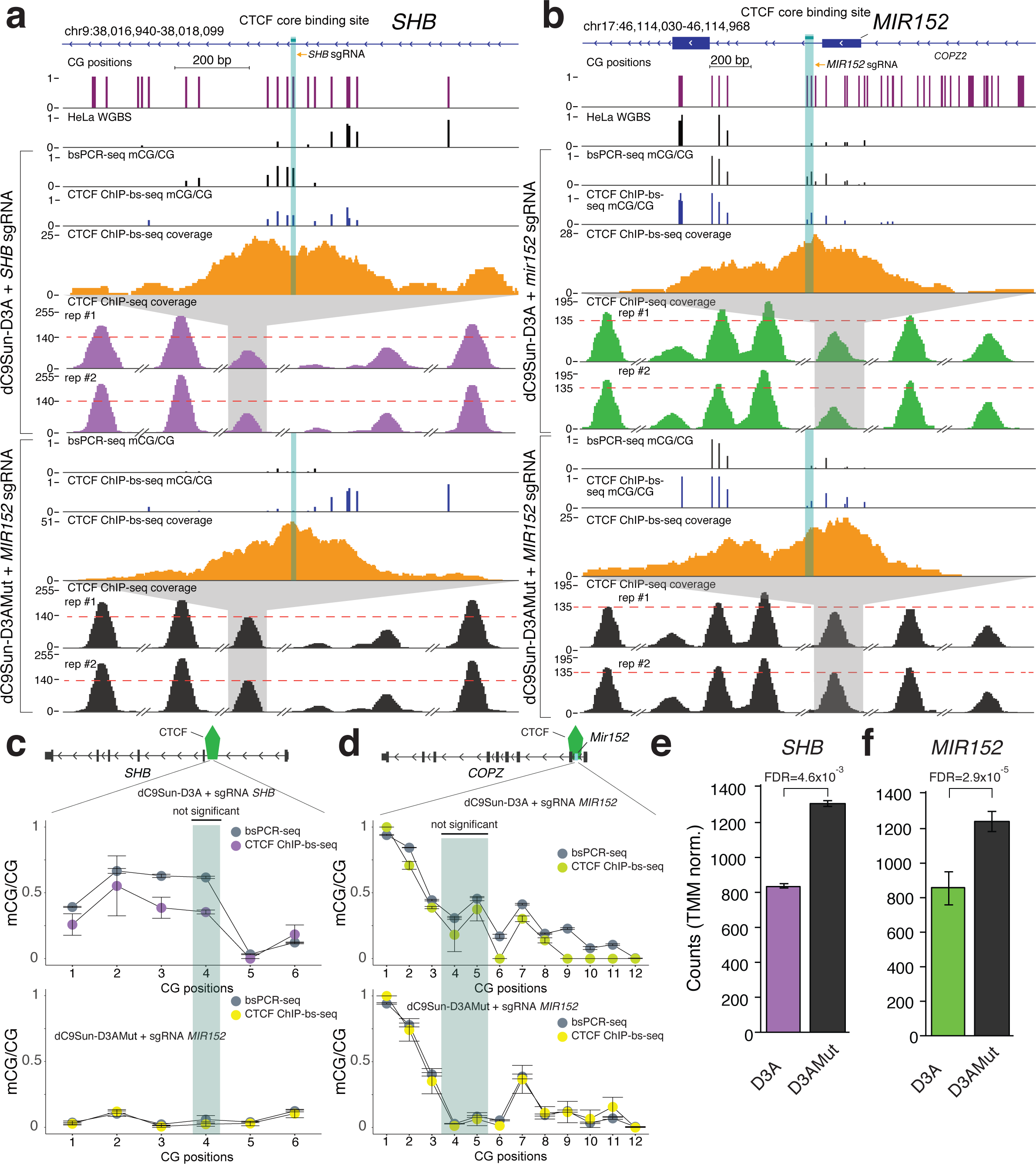
Impact of targeted DNA methylation induction on CTCF binding. **(a)** Genome Browser display of targeted CTCF binding site in SHB intron. Sets of experiments include from top to bottom: targeted bsPCR-seq (fraction mCG/CG), CTCF-ChIP-bs-seq (fraction mCG/CG), CTCF-ChIP-bs-seq coverage and CTCF ChIP-seq coverage. CTCF core binding site is highlighted in shaded green. CTCF ChIP-seq coverage (TMM normalized counts) are shown with adjacent most peaks for reference (broken x-axis). Red dotted line is set to maximum targeted CTCF peak in the control samples. **(b)** Genome browser snapshot of targeted CTCF binding site upstream of *MIR152*. **(c)** and **(d)** Quantitation of mCG/CG in CTCF core binding site (green shaded region) and adjacent to core binding site comparing targeted bsPCR-seq (gray circles) to ChIP-bs-seq (purple or green circles for aGCN4-D3A and yellow circles for aGCN4-D3AMut) for *SHB* and *MIR152*, respectively (replicates n=2, sd - error bars, statistic: Fisher’s exact test. Quantitation of CTCF TMM normalized ChIP-seq peak at *SHB* **(e)** and *MIR152* **(f)**, respectively (replicates n=2, sd - error bars, statistic edgeR, Benjamini-Hochberg multiple test corrected p-values)).

To to extend the findings of targeted mCG deposition on CTCF binding to an additional TF, we next investigated the impact of mCG deposition on the binding of NRF1. This TF was selected as previous studies have found that NRF1 binding appears to be DNA methylation sensitive *in vitro*, as determined by a differential array binding assay (Hu et al. 2013), as well as in cell culture, where NRF1 occupies binding sites that lose CG methylation in *DNMT* triple knockout embryonic stem cells. (Domcke et al. 2015). Similarly to our CTCF analyses, we set out to target four NRF1 occupied binding sites in HeLa cells, with each having at least one hypomethylated CG in the core binding site (*TMEM206* promoter, downstream of *TRAPPC3*, upstream of *MSANTD3*, and downstream of *TEF*) with six different sgRNAs each. These targets were chosen based on both strong NRF1 occupancy as well as having narrow hypomethylated binding sites, where methylation was found within a 1 kb window both upstream and downstream of the NRF1 binding site. As noted before (Hinz et al. 2015; Horlbeck et al. 2016; Moreno-Mateos et al. 2015), and seen in the resulting on-target DNA methylation patterns, sgRNA placement greatly affected the extent of CG methylation induced by dC9Sun-D3A (Supplementary Fig. 12a-d). Consequently, we focussed was solely on the *TMEM206* promoter as it showed the greatest susceptibility to targeted mCG induction based on the presence of three CGs in its 15 bp core NRF1 binding site, and appeared to be strongly occupied by NRF1 (Fig. 5a). To maximize on-target DNA methylation deposition for all three CGs, we chose sgRNA #2 for all further experiments (Supplementary Fig. 12a). Next we performed targeted bsPCR-seq to measure the average DNA methylation induction for all puromycin selected cells, and found a 30%, 32% and 32% increase in methylation level for each of the three CGs in the *TMEM206* core binding site, respectively (Fig. 5b). As with our CTCF analyses, we then tested

**Figure 5.**
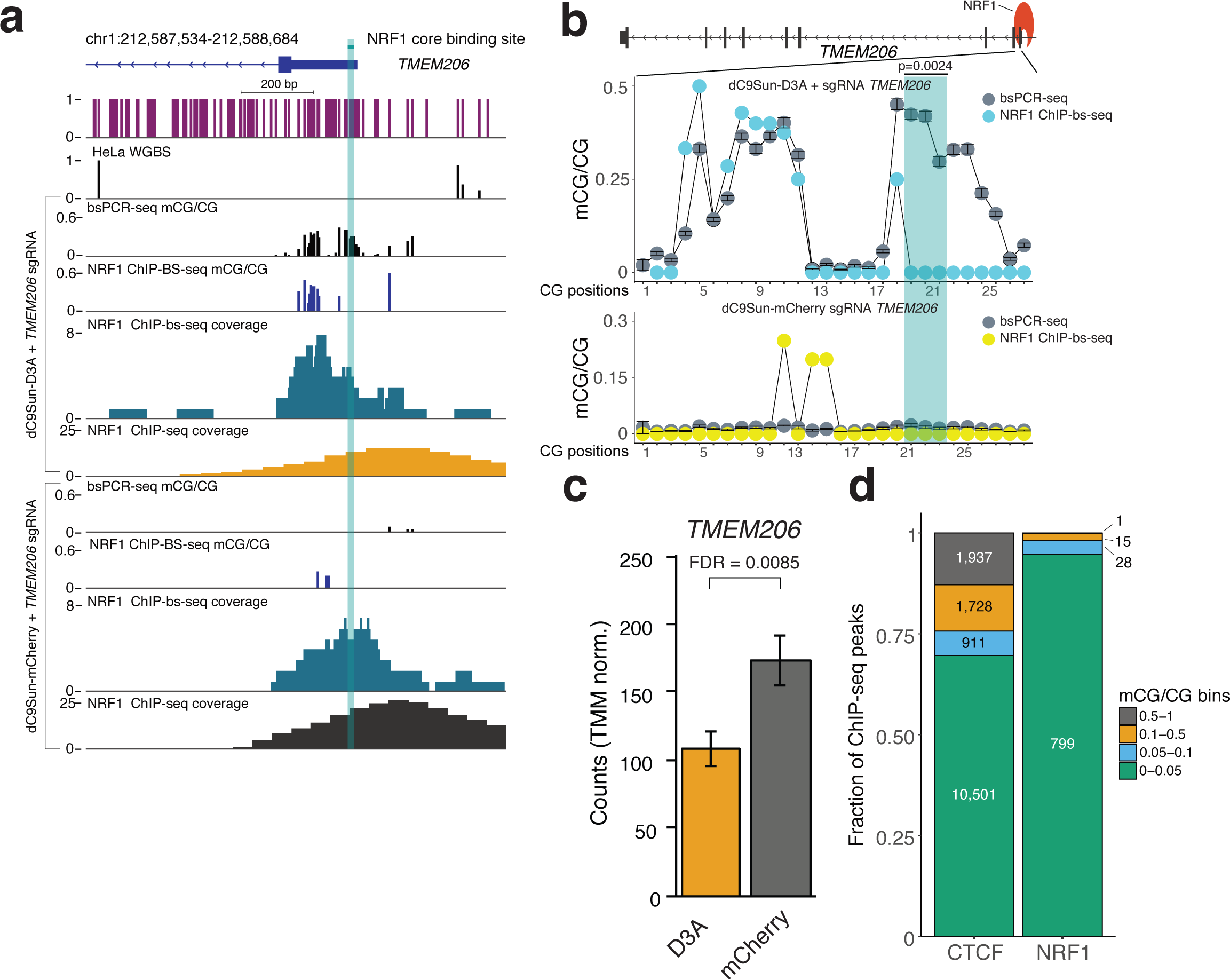
Impact of targeted DNA methylation induction on NRF1 binding. **(a)** Genome Browser display of targeted NRF1 binding site in TMEM206 promoter. Sets of experiments include from top to bottom bsPCR-seq (mCG/CG), NRF1-ChIP-bs-seq (mCG/CG), NRF1-ChIP-bs-seq coverage and NRF1 ChIP-seq-coverage. NRF1 core binding site is highlighted in shaded green. **(b)** Quantitation of mCG/CG in NRF1 core binding site (green shaded region) and adjacent to core binding site, comparing targeted bsPCR-seq (gray circles) to ChIP-bs-seq (cyan circles for aGCN4-DNMT3A and yellow circles for aGCN4-mCherry) at the *TMEM206* promoter (NRF1 ChIP-bs samples combined n=3 for coverage, bsPCR-seq, n=3, sd - error bars, statistic: Fisher’s exact test). **(c)** Quantitation of TMEM206 promoter NRF1 ChIP-seq peak counts in samples treated with aGCN4-DNMT3aWT (orange) compared to aGCN4-mCherry (gray). Peak counts were TMM normalized with THOR program (mCherry n=2, D3A n=3, statistic edgeR, Benjamini-Hochberg multiple test corrected p-values). **(d)** Comparison of average mCG/CG in CTCF and NRF1 core binding sites, respectively by binning them into intervals of no (≥ 0 and ≤ 0.05), low (> 0.05 and ≤ 0.1), intermediate (> 0.1 and ≤ 0.5) or high (> 0.5) levels of DNA methylation.

whether NRF1 could also bind to methylated DNA at the *TMEM206* promoter by performing NRF1 ChIP-bs-seq. Due to the very low quantity of NRF1-immunoprecipitated DNA as the result of fewer NRF1 binding sites in the genome as compared to other DNA binding proteins such as CTCF, we merged data from multiple replicates (n=3) to gain sufficient coverage to accurately quantitate mCG levels in the NRF1 core binding site. The NRF1 core binding site and CGs up- and downstream of it are completely hypomethylated as judged by bsPCR-seq and NRF1 ChIP-bs-seq when dC9Sun-mCherry is recruited to the *TMEM206* promoter (Fig 5b, lower panel). Interestingly, when *TMEM206* promoter was targeted by dC9Sun-D3A, no mCG was detectable in the ChIP-bs-seq data of the NRF1 core binding site (Fig. 5b), however upstream of this binding site mCG was still detectable to the same level as seen previously in the targeted bsPCR-seq (Fig. 5b). This observation suggested that mCG was only tolerated upstream of the NRF1 binding site. Consequently, NRF1 either blocked mCG deposition in its core binding site or lost its binding capacity once mCG was deposited by dC9Sun-D3A. To understand the impact on NRF1 binding, we performed NRF1 ChIP-seq and found that NRF1 binding is reduced by 30% (edgeR, FDR=0.0085, ranked 63 out of 607) when targeted by dC9Sun-D3A compared to cells targeted by dC9Sun-mCherry (Fig. 5c). Furthermore, while dC9Sun-D3A may have the potential to recruit other factors that cause the observed reduction in NRF1 binding, this would not be consistent with the observed DNA methylation pattern of NRF1-bound DNA as measured by ChIP-bs-seq (Fig. 5b).

To compare the different levels of CTCF and NRF1 DNA methylation tolerance, core binding sites in the genome were interrogated by extracting the position of CGs in each core binding site (CTCF and NRF1 positional weight matrix) followed by averaging mCG/CG levels at all CG sites within each core binding site of CTCF and NRF1 ChIP-bs-seq data. Further, we parsed the average level of mCG/CG per core binding site for CTCF and NRF1 into bins based on the average mCG level (Fig. 5d, TF binding motif mCG level <5%, 5%-10%, 10%-50%, and >50%), where 94.8% (799 out of 843) of NRF1 binding sites genome-wide have DNA methylation levels less than 5%. In contrast, only 69.6% (10,501 out of 15,077) of CTCF binding sites have DNA methylation levels less than 5%, which may explain why we detected that CTCF could bind to it’s core binding site even after targeted DNA methylation induction by dC9Sun-D3A (Fig. 4 c, d). Taken together, we show that targeted DNA methylation, but not dCas9 interference (Fig. 4 c, d, h), displaced both CTCF and NRF1, with the latter showing the higher DNA methylation sensitivity. Future work could leverage precise mCG deposition of the dC9Sun-D3A system to test direct mCG sensitivity for a variety of TFs *in vivo*.

## Discussion

Previous work has highlighted the ability of dCas9 to be used to induce DNA methylation at desired regions in the genome(Vojta et al. 2016; Amabile et al. 2016; Liu et al. 2016; Lei et al. 2017; Stepper et al. 2017). However, a key issue that to date has not been sufficiently addressed is the potential for off-target DNA methylation induction, which could lead to non-specific genomic responses and limits the utility of the systems for unambiguous assessment of the effect of DNA methylation at a locus. Single fusion constructs, such as dC9-D3A, lack the ability to fine-tune on- versus off-target DNA methylation deposition, and exhibit extensive off-target activity. When dC9-D3A expression is increased to attain high on-target methylation, the off-target methylation rate reached unacceptably high levels. Conversely, limiting the expression of the single fusion to reduce off-target methylation levels resulted in poor induction of on-target DNA methylation. Therefore, the single fusion constructs present multiple significant deficiencies. Here, we present a highly specific and tunable system to perform targeted alteration of DNA methylation, based on the modular SunTag system(Tanenbaum et al. 2014), allowing independent variation of the expression of the DNA targeting module (dC9Sun) and the effector module (αGCN4-D3A). While no significant difference in the distance at which DNA methylation deposition was observed between the dC9Sun-D3A system and the direct fusion systems, dC9Sun-D3A exhibited greatly improved specificity for depositing targeted methylation compared to dC9-D3A, and achieved effective targeting of multiple loci simultaneously with on-target DNA methylation induction levels comparable to single-guide experiments. Recently, a similar dC9Sun system employing full length DNMT3A1 was described by Huang *et al.(Huang et al. 2017)*, where lentiviral delivery was used. Huang *et al.* investigated off-target mCG deposition by reduced representation bisulfite sequencing (RRBS) at a limited subset of CpG islands (∼6100, 21.3% of all possible CpG islands), with a single sample per condition and reported off-target mCG deposition to be lower than dC9-D3A direct fusion systems. In contrast, we employed the more comprehensive approach of using TSC-bs-seq, covering 2.1 million CGs at >= 5 reads per sample (∼11x the number of CGs surveyed in Huang *et al* at equivalent coverage). The Illumina EPIC TCS-bs-seq system is designed to cover a large number of putative regulatory regions in the genome that are frequently in an unmethylated state, here covering ∼12.7% of all hypomethylated CGs (at >= 5 reads per condition) in the HeLa genome (∼5.5×10^6^ hypomethylated CGs in HeLa genome, mCG/CG <= 0.2). In contrast, the method used by Huang *et al.* would only cover ∼1.8% of the hypomethylated CG sites in the HeLa genome. We found that our system was ∼40-fold less susceptible to gain or loss of mCG at CG islands compared to the system described by Huang *et al.* (Supplementary Fig. 13). Thus, through well replicated use of this TCS-bs-seq system, we have performed the most extensive survey of on- and off-target methylation induced by dCas9-based DNA methylation modifying systems to date, demonstrating that our dC9Sun-D3A system is highly specific.

Having established the specificity and fidelity of our dC9Sun-D3A system, we subsequently directly determined the impact of targeted DNA methylation on the binding of two DNA binding proteins, CTCF and NRF1. Although CTCF sensitivity to DNA methylation had previously been reported(Liu et al. 2016),(Lei et al. 2017), our work is the first to directly test the impact of targeted DNA methylation deposition upon NRF1 binding. Our results indicate that both CTCF and NRF1 are impacted by DNA methylation deposition. Surprisingly, CTCF was still able to tolerate and bind methylated DNA in its core binding site, for three independent sites. A possible explanation for this is that the binding of CTCF is in a transitional state, where its core binding site has been methylated but CTCF has yet to be evicted. Alternatively, CTCF may be binding 5-hydroxymethylcytosine (5hmC), the byproduct of Tet mediated oxidation of 5-methylcytosine, 5mC), as previously described(Marina et al. 2016; Teif et al. 2014; Feldmann et al. 2013) since CTCF ChIP-bs-seq cannot distinguish between 5mC and 5hmC. In contrast to CTCF, we did not observe NRF1 binding to any intermediate 5mC or possible 5hmC states, as there was a loss in NRF1 occupancy upon DNA methylation deposition. Taken together, the dC9Sun-D3A technology opens up avenues to perform highly precise DNA methylation deposition and could be adapted to effectively implement alternative epigenetic editing such as altering histone modifications or DNA demethylation with the same advantages as described above. It is also conceivable to leverage the array of GCN4 binding sites on the SunTag for multiplexed epigenetic editing, employing more than one epigenetic effector simultaneously.

## Conclusion

The influence of DNA methylation changes on gene regulation is still a highly debated topic, with broad implications for interpretation of the potential effect of differential methylation states evident in a variety of contexts, such as development and cancer. However, it is paramount to study the effects of targeted DNA methylation changes at key regions with a high degree of confidence, such that off-target effects by spurious DNA methylation deposition are minimized. To that end, the dC9Sun-D3A system described here is highly adaptable and tunable, and capable of high on-target mCG deposition while not suffering from spurious off-target DNA methylation induction as observed with some other previously described dCas9 systems. Therefore, the dC9Sun-D3A system could be employed in a variety of different cell types and systems to test the direct effect of mCG deposition on TF binding, splicing or transcription in general.

## Methods

### Cell lines

MCF-7 and HeLa cells were cultured in a humidified cell culture incubator at 37^°^C with 5% (v/v) CO_2_ as previously described (Rhee et al., 2002). MCF-7 cells were maintained using MEM (Minimum Essential medium) alpha (Life technologies, Cat no: 12571071) supplemented with 10% (v/v) fetal bovine serum (FBS) (Integrated Sciences, Cat no: HYCSV3017603), 1X (v/v) 5.5% Sodium bicarbonate (Life technologies, Cat no: 25080094) and 1X (v/v) Glutamax (Life technologies, Cat no: 35050061) while HeLa cells were cultured with DMEM (Dulbecco’s Modified Eagle Medium) supplemented with 10% (v/v) FBS and 1X (v/v) Glutamax.

### Cell transfections

Transfections were carried out using a 1:3 ratio of DNA to FuGENE HD (Promega, Cat no: E2311) following manufacturer’s instructions. Briefly, MCF-7 or HeLa cells were seeded at 80% confluency in either 6-wells culture plate (In-vitro Technologies, Cat no: FAL353046) or 15 cm culture dishes (In-vitro Technologies, Cat no: COR430599). 24 hours post-seeding, 500 ng of TALE-DNMT3A or dCas9-DNMT3A constructs and 500 ng of individual sgRNA plasmids were transfected into cells. For the TALE and dCas9-SunTag constructs, 330 ng was used, along with 330 ng of individual or pooled sgRNAs, 40 ng of αGCN4-DNMT3A, and 300-320 ng of pEF1a-GFP-Puro (transfection control plasmid) to make the final amount to 1 ug. This was scaled up for 15 cm plate transfections. Cells were screened after 48 hours for expression of reporter gene (GFP or mCherry) by fluorescent microscopy followed by either FACS sorting or 48 hours of puromycin selection (2 µg/mL, Life technologies, Cat no: A1113803) to enrich for positively transfected cells. After puromycin selection, surviving cells were subsequently either used for chromatin immunoprecipitations, chromatin immunoprecipitation followed by bisulfite sequencing or for targeted bisulfite sequencing where DNA was extracted by the ISOLATE II DNA extraction kit (Bioline, Cat no: BIO-52067). Triple positive cells containing the tagBFP2 (dCas9-Suntag vector), mCherry (sgRNA vector) and sfGFP(αGCN4-DNMT3A vector) were FACS sorted and subjected to DNA extraction using the ISOLATE II DNA extraction kit (Bioline, Cat no: BIO-52067) which was further used in determining genome-wide off-target DNA methylation deposition using the TruSeq Methyl Capture EPIC library prep kit (Illumina, Cat no: FC-151-1002) as well as for targeted bisulfite sequencing.

### Plasmid construction

The human codon optimized Cas9 was amplified using PCR from the hCas9 D10A plasmid (Addgene, 41816). Subsequently, PCR mutagenesis was conducted to introduce the H840A mutation to generate the catalytically dead Cas9 (dCas9). The DNMT3A sequence was PCR amplified from the ZF-598-DNMT3A plasmid (Gift from Pilar Blancafort) and the N-terminal 3xHA tag, linker peptide between dCas9 and DNMT3A, and C-terminal 3xTy1 tag was ordered as separate gBlocks (IDT). Finally, the dCas9 cassette was inserted into a plasmid backbone containing a puromycin selectable marker. The single fusion TALE expression vector backbone was based on (Sanjana et al. 2012), and Gibson assembly (Gibson et al. 2009) was used to replace the nuclease with DNMT3A. The linker peptide for the TALE-DNMT3A expression vector was identical to the dCas9 counterpart.

Gibson Assembly was used to clone the empty TALE-SunTag backbone using plasmids pHRdSV40-dCas9-10XGCN4_v4_P2A-BFP (Addgene, 60903) and pEF1a-TALE-DNMT3A-WT (Addgene, 100937). Similarly, the dCas9-SunTag construct was assembled from pGK-dCas9-DNMT3A-WT (V3; Addgene, 100938), pHRdSV40-dCas9-10XGCN4_v4_P2A-BFP (Addgene, 60903) and a polyA gBlock sequence (IDT). Following this, pHRdSV40-scFv-GCN4-sfGFP-VP64-GB1-NLS (Addgene, 60904), pEF1a-TALE-DNMT3A-WT (Addgene, 100937) and a gBlock linker sequence (IDT) was used to assemble αGCN4-DNMT3A, which was also used to create the DNMT3A FEER>AAAA mutation. This same backbone was used to Gibson assemble the αGCN4-mCherry control plasmid, where the DNMT3A domain was replaced with a mCherry fluorophore. Furthermore, restriction ligation was used to insert TALE targeting sequences within the empty TALE-SunTag backbone using MluI and AfeI (NEB). TALE DNA binding domain inserts were synthesised in-house using Iterative Capped Assembly as previously described (Briggs et al. 2012). Finally, gRNA plasmids were synthesised according to the protocol by (Mali et al. 2013) and target sequences for TALEs and gRNA are listed in Supplementary Table 2. All plasmids constructed for this study can be found here: https://www.addgene.org/plasmids/articles/28191639/.

### Targeted BS-PCR primer design and sequencing

For each bisulfite PCR amplicon, primers were designed with MethPrimer tool (http://www.urogene.org/cgibin/methprimer/methprimer.cgi) with all parameters set to default except the following: product size 200-400 bp; primer Tm min: 55°C, opt: 60°C and max: 64°C; primer size min: 20 nt, opt: 25 nt and max: 30 nt. Each primer pair was tested in a 15 µL PCR reaction with either NEB EpiMark (NEB, M0490L) or 2X MyTaq (Bioline, BIO-25041) for a range of annealing temperatures ranging from 50°C to 64°C for 40 cycles. Primers that amplified specific products of the expected size were used and these amplicons were confirmed as our regions of interest via sanger sequencing. 500ng extracted genomic DNA from transfected HeLa or MCF-7 cells and 0.5% spiked in lambda DNA (non-conversion control) was bisulfite converted using the EZ DNA-Methylation-Gold kit (Zymo Research, Cat no: D5006) following the manufacturer’s instructions. 20 ng of bisulfite converted DNA was used for generating PCR products, which were subsequently pooled in an equimolar ratio and purified by Agencourt AMPure XP beads (Beckman Coulter, Cat. No: A63880). Libraries were prepared by A-tailing and ligating Illumina-compatible Y-shaped adapters to amplicons which were amplified by three PCR cycles and subsequently purified by Agencourt AMPure XP beads (Beckman Coulter, Cat no: A63880). Single-end 300 bp sequencing was performed on an Illumina MiSeq platform.

### ChIP-sequencing

Transfected HeLa cells were used for performing chromatin immunoprecipitations for CTCF and NRF1 as described previously (Wang et al. 2012) (Domcke et al. 2015) with some modifications. Briefly, cells were crosslinked for 10 minutes in 1% formaldehyde and quenched in 125 mM glycine. Chromatin was sheared (Covaris, S220) for 5 min, 5% duty cycle, 200 cycles per burst and 140 watts peak output and incubated with either a CTCF (Cell Signaling, 2899B), NRF1 (Abcam, ab55744), HA (Biolegend, 901502) or Ty1 (Sigma, SAB4800032) antibody and subsequently conjugated to a 50:50 mix of protein A/G Dynabeads (Invitrogen, M-280). After wash steps, DNA was eluted, crosslinks were reversed, and immunoprecipitated DNA was purified by Agencourt AMPure XP beads (Beckman Coulter, Cat no: A63880). Libraries were prepared from the entire ChIP eluate volume containing either 1ng or 0.1ng input material per replicate for CTCF or NRF1, respectively, using the ThruPLEX DNA-seq 12S kit (Rubicon Genomics, R400429). After limited PCR amplification, libraries were purified using Agencourt AMPure XP beads (Beckman Coulter, Cat no: A63880), and eluted in a final volume of 20 ul. Single-end 100 bp sequencing was performed on a HiSeq 1500.

### Chromatin immunoprecipitation bisulfite sequencing

Chromatin immunoprecipitation was performed using CTCF and NRF1 antibodies as stated above. Either 1ng or 10ng per replicate of CTCF and NRF1 ChIP eluate was used and subjected to bisulfite conversion using Zymo methylation direct kit (Zymo Research, D5020). Bisulfite converted DNA was immediately handled for library preparation using the Accel-NGS Methyl-Seq DNA library kit (Swift Biosciences, Cat no: 30024) following the manufacturer’s instructions. Libraries were amplified for either 9 or 13 cycles for CTCF and NRF1, respectively. Single-end 100 bp sequencing was performed on a HiSeq 1500.

### Targeted enrichment for determining off-target DNA methylation deposition

Hela cells were transfected with pEF1a-GFP-Puro and pEF1a-mCherry for control baseline or pGK-dCas9-Suntag-tagBFP2, pEF1a-mCherry-SHB1 sgRNA and pEF1a-aGCN4-DNMT3A for targeted methylation at *SHB* locus in triplicate. Transfected cells were sorted for GFP and mCherry positive (control baseline) or GFP, mCherry and BFP positive (targeted methylation at *SHB* locus) and subjected to DNA extraction using the ISOLATE II DNA extraction kit (Bioline, Cat no: BIO-52067). 250 ng of genomic DNA from each replicate was used for library preparation and subsequently pooled for target enrichment using the TruSeq Methyl Capture EPIC library prep kit (Illumina, Cat no: FC-151-1002) following manufacturer’s instructions. Libraries were bisulfite converted using EZ DNA methylation Lightning kit (Zymo Research, D5030) and amplified for 10 cycles. Single-end 100 bp sequencing was performed on a HiSeq 1500.

### Western blot

HeLa cells were seeded at 80% confluency and transfected with equal amounts of sterilized plasmids dC9-D3A high (or dC9-D3A) and SHB-1-sgRNA to 15.8 µg, along with three times the amount of FuGENE HD transfection reagent (Promega, Cat no: E2311) following manufacturer’s instructions. Samples were either collected 48 h post-transfection or following an additional 48 h after Puromycin selection (2 µg/mL, Life technologies, Cat no: A1113803) to replicate the conditions employed while generating cells used for DNA methylation analysis. Cells were harvested and spun down at 300 x g for 5 minutes, then lysed in RIPA buffer with 4% SDS and 1 µl Benzonase (Sigma, Cat no: E1014). Samples were further lysed by sonication (Covaris, S220) and gently pipetted with a 21G needle (Livingston, Cat no: DN21Gx1.0LV) to reduce viscosity. Standard immunoblotting was performed on 50 µg of cell lysate for each sample. Mouse anti-Ty1 (1:1,000, Sigma, Cat no: AB4800032) antibody was used, along with mouse anti-α-Tubulin (1:1,000, GenScript, Cat no. A01410-40) as the loading control.

### Data processing

ChIP-Seq and ChIP-bs-seq data was quality checked and hard trimmed (Accel-NGS Methyl-Seq DNA library kit requirements) using Trimmomatic (Bolger et al. 2014) with the following parameters: ILLUMINACLIP:2:30:10, LEADING:3, TRAILING:3, SLIDINGWINDOW:4:20, MINLEN:25. Trimmed ChIP-seq data were mapped to the human genome (GRCh37/hg19) using Bowtie1.1.1 (Langmead et al., 2009) with the options ‘‑‑wrapper basic-0 -m 1 -S -p 4 -n 1’’ allowing up to one mismatch. Reads mapping to multiple locations were then excluded, and reads with identical 5’ ends and strand were presumed to be PCR duplicates and were excluded using Picard MarkDuplicates. Bigwig coverage tracks and differential peak calling were produced by THOR (Allhoff et al. 2016) using TMM normalization. BS-PCR-Seq data and ChIP-bs-seq data was mapped to the GRCh37/hg19 genome and processed using BS-Seeker2 (Guo et al. 2013) with default parameters, Bowtie 1 and allowing for one mismatch.

### Availability of data and materials

Raw and processed data of ChIP sequencing, bsPCR bisulfite sequencing and ChIP bisulfite sequencing have been deposited to the Gene Expression Omnibus (GSE107607).

## Funding

This work was funded by the Australian National Health and Medical Research Council (GNT1069830), the Australian Research Council (ARC) Centre of Excellence program in Plant Energy Biology (CE140100008), the National Institutes of Health (5R01DA036906), and the Raine Medical Research Foundation. Support was provided by an ARC Future Fellowship to RL (FT120100862), Sylvia and Charles Viertel Senior Medical Research Fellowships (to Rl and JMP), and a Howard Hughes Medical Institute International Research Scholarship to RL.

## Acknowledgments

The authors thank O. Bogdanović, P. Blancafort and the members of the Lister and Polo laboratories for helpful discussions and feedback.

## Ethics approval

Not applicable in this study.

## Competing interests

The authors declare that they have no competing interests

